# Muscle mechanics and energy expenditure of the triceps surae during rearfoot and forefoot running

**DOI:** 10.1101/424853

**Authors:** Allison H. Gruber, Brian R. Umberger, Ross H. Miller, Joseph Hamill

## Abstract

Forefoot running is advocated to improve running economy because of increased elastic energy storage than rearfoot running. This claim has not been assessed with methods that predict the elastic energy contribution to positive work or estimate muscle metabolic cost. The purpose of this study was to compare the mechanical work and metabolic cost of the gastrocnemius and soleus between rearfoot and forefoot running. Seventeen rearfoot and seventeen forefoot runners ran over-ground with their habitual footfall pattern (3.33-3.68m•s^−1^) while collecting motion capture and ground reaction force data. Ankle and knee joint angles and ankle joint moments served as inputs into a musculoskeletal model that calculated the mechanical work and metabolic energy expenditure of each muscle using Hill-based muscle models with contractile (CE) and series elastic (SEE) elements. A mixed-factor ANOVA assessed the difference between footfall patterns and groups (α=0.05). Forefoot running resulted in greater SEE mechanical work in the gastrocnemius than rearfoot running but no differences were found in CE mechanical work or CE metabolic energy expenditure. Forefoot running resulted in greater soleus SEE and CE mechanical work and CE metabolic energy expenditure than rearfoot running. The metabolic cost associated with greater CE velocity, force production, and activation during forefoot running may outweigh any metabolic energy savings associated with greater SEE mechanical work. Therefore, there was no energetic benefit at the triceps surae for one footfall pattern or the other. The complex CE-SEE interactions must be considered when assessing muscle metabolic cost, not just the amount of SEE strain energy.

## Introduction

Effective storage and release of elastic strain energy within muscles, particularly the triceps surae group, is suspected to contribute to efficient locomotion in humans and other animals (e.g., Cavagna et al., 1977b; Ishikawa et al., 2007). The triceps surae (gastrocnemius and soleus) has been extensively studied because of its potential for storage and release of elastic energy during the stance phase of gait and because it is a major contributor to mechanical work and the support moment in locomotion (Alexander and Bennet-Clark, 1977; Biewener and Roberts, 2000; Sasaki and Neptune, 2006; Winter, 1983). In human running, the mid- and forefoot running patterns, characterized by initial ground contact on the anterior portion of the foot, have been suggested to store and return more triceps surae elastic strain energy per stride than the common rearfoot pattern with initial ground contact by the heel (Hasegawa et al., 2007; Hof et al., 2002; Lieberman et al., 2010; Perl et al., 2012). However, the hypothesis that the use of forefoot patterns rather than a rearfoot pattern should lead to more efficient running has primarily received indirect support from evidence that most elite human runners use the forefoot pattern when racing (Kerr et al., 1983; Payne, 1983).

In light of these suspected whole-body energetic benefits of the forefoot pattern, it is perhaps surprising that most humans (~75%) use a rearfoot pattern when running (e.g., Hasegawa et al., 2007; Larson et al., 2011). Although humans have numerous physiological features that seem to enable especially good endurance running on a comparative basis (Bramble and Lieberman, 2004), our net mass-specific whole-body metabolic cost for running is on average slightly greater than expected for an animal of our body size (Rubenson et al., 2007). Perhaps our preference for the rearfoot pattern causes this disparity. For example, it has been argued that the rearfoot pattern limits the whole-body running economy and the storage and return of elastic strain energy compared with rearfoot running (Ardigo et al., 1995; Bramble and Lieberman, 2004; Perl et al., 2012; Schmitt and Larson, 1995). However, studies on human running energetics with different footfall patterns have failed to observe reduced rates of oxygen consumption with the forefoot pattern than with the rearfoot pattern (Ardigo et al., 1995; Cunningham et al., 2010; Gruber et al., 2013; Lussiana et al., 2017; Ogueta-Alday et al., 2013; Perl et al., 2012). Given that these findings conflict with the suggestion that forefoot patterns should result in a reduced whole-body metabolic cost, greater elastic energy storage and release by the triceps surae may not be a mechanism that contributes significantly to metabolic energy savings in the forefoot versus the rearfoot pattern in human running.

Differences in muscle-level energetics between rearfoot and forefoot running have yet to be compared directly. Earlier studies have observed that forefoot running results in greater Achilles tendon forces during the stance phase that may contribute to greater elastic energy storage and release than rearfoot running (Almonroeder et al., 2013; Perl et al., 2012). However, given that recent findings have indicated no difference in running economy between habitual rearfoot and forefoot runners, nor an immediate improvement in running economy by changing between rearfoot and forefoot running (Ardigo et al., 1995; Cunningham et al., 2010; Gruber et al., 2013; Lussiana et al., 2017; Ogueta-Alday et al., 2013; Perl et al., 2012), it may be that even if there is greater storage and release of elastic energy in forefoot running it does not result in any whole-body metabolic energy savings compared with rearfoot running. For example, greater force generation will increase the tendon stretch and thus the potential for greater energy storage and more economical running by allowing the muscle fibers to act more isometrically, but greater muscle force can also increase the metabolic energy consumption of the muscle because of increased activation (Biewener and Roberts, 2000; Roberts et al., 1998).

Previous studies comparing footfall patterns have been largely based on inverse dynamics analyses of resultant ankle joint moments during the stance phase. Inverse dynamics analyses do not directly account for individual muscle forces, fiber-tendon interactions, and contributions of individual muscles to mechanical energetics or metabolic energy expenditure in locomotion (Sasaki et al., 2009; Zajac et al., 2002; Zajac et al., 2003). Direct measurements of individual muscle mechanics and muscle energetics are ideal methods for investigating muscle function (Umberger and Rubenson, 2011) but are impractical in humans because they require invasive surgical procedures and specialized equipment. Hill-based muscle models provide a noninvasive approach for estimating triceps surae muscle function in human locomotion (Erdemir et al., 2007; Hof et al., 2002). Studies modeling the triceps surae during human locomotion have successfully predicted muscle-tendon interactions and have been able to distinguish differences in muscle component length changes, forces, and mechanical work between individuals and conditions (e.g., Farris and Sawicki, 2012; Fukunaga et al., 2001; Scholz et al., 2008).

The purpose of this study was to compare the muscle mechanical work and muscle metabolic energy expenditure of the triceps surae muscle group between rearfoot and forefoot running patterns using experimental gait analysis combined with Hill-based muscle modeling. Given the indication that forefoot running results in greater Achilles tendon forces, forefoot running may result in greater tendon stretch and elastic energy storage. This greater elastic energy return during the stance phase of forefoot running can potentially reduce muscle metabolic energy expenditure if it results in the contractile elements operating at more optimal contraction velocities compared with rearfoot running (Alexander, 2002; Biewener and Roberts, 2000). It was then hypothesized that rearfoot running would result in greater contractile element mechanical work from the triceps surae muscles, whereas forefoot running would result in greater series elastic element mechanical work (i.e., greater storage and release of elastic energy) from the triceps surae muscles. Secondly, as a result of a greater contribution from elastic energy and less contractile element mechanical work, it was hypothesized that forefoot running would result in lower muscle metabolic energy expenditure than rearfoot running.

## Materials and Methods

### Participants

Thirty-four experienced, habitual rearfoot and forefoot runners from the local community were included in this study. Participants were included if they ran at least 16 km/week (mean±SD: 42.1±29.0 km/week), were free of any cardiovascular or neurological disorders, and had not experienced any injuries, surgeries, or other impairments that affected their running gait in the past year. Participants had an average preferred endurance running speed of 3.7±0.3 m•s^−1^ for leisurely runs. All participants read and completed an informed consent document and questionnaires approved by the university Institutional Review Board before participating.

### Experimental Setup

Participants wore form-fitting shorts and tops and neutral, lightweight “racing flat” shoes provided by the laboratory (RC 550, New Balance, Brighton, MA, USA, 215.7 g). A set of spherical retro-reflective markers were attached to the pelvis and right thigh, shank, and shoe over anatomical landmarks of the foot according to established methods (Hamill et al., 2014).

Marker positions were sampled at 240 Hz using an eight-camera optical motion capture system (Oqus 300, Qualisys, Gothenburg, Sweden). Ground reaction forces (GRF) were sampled synchronously with the marker positions at 1200 Hz using a large force platform (OR6-5, AMTI, Watertown, MA, USA). The force platform was embedded flush with the floor at the center of a level 25-m runway and surrounded by the motion capture cameras. Running speed was determined using photoelectric sensors (Lafayette Instruments, Lafayette, IN, USA) positioned 3-m before and after the center of the force platform.

### Determination of Habitual Footfall Pattern and Groups

Habitual footfall patterns (rearfoot and forefoot) were determined for each participant by recording high-speed digital video (Exilim EX-F1, Casio Computer Co., LTD, Shibuya-ku, Tokyo, Japan) sampling at 300 Hz while the participants ran over-ground at their preferred endurance running speed. The starting position was adjusted so that the participants landed with the right foot in the center of the force platform. Participants were classified into the habitual forefoot group if they had a strike index greater than 34% (Cavanagh and Lafortune, 1980), a diminished or absent vertical impact peak in their GRF profile (Figure 1), and an ankle with 0° plantar flexion or more at initial contact. Participants not meeting these criteria were included in the rearfoot group (Table 1).

**Figure 1:**
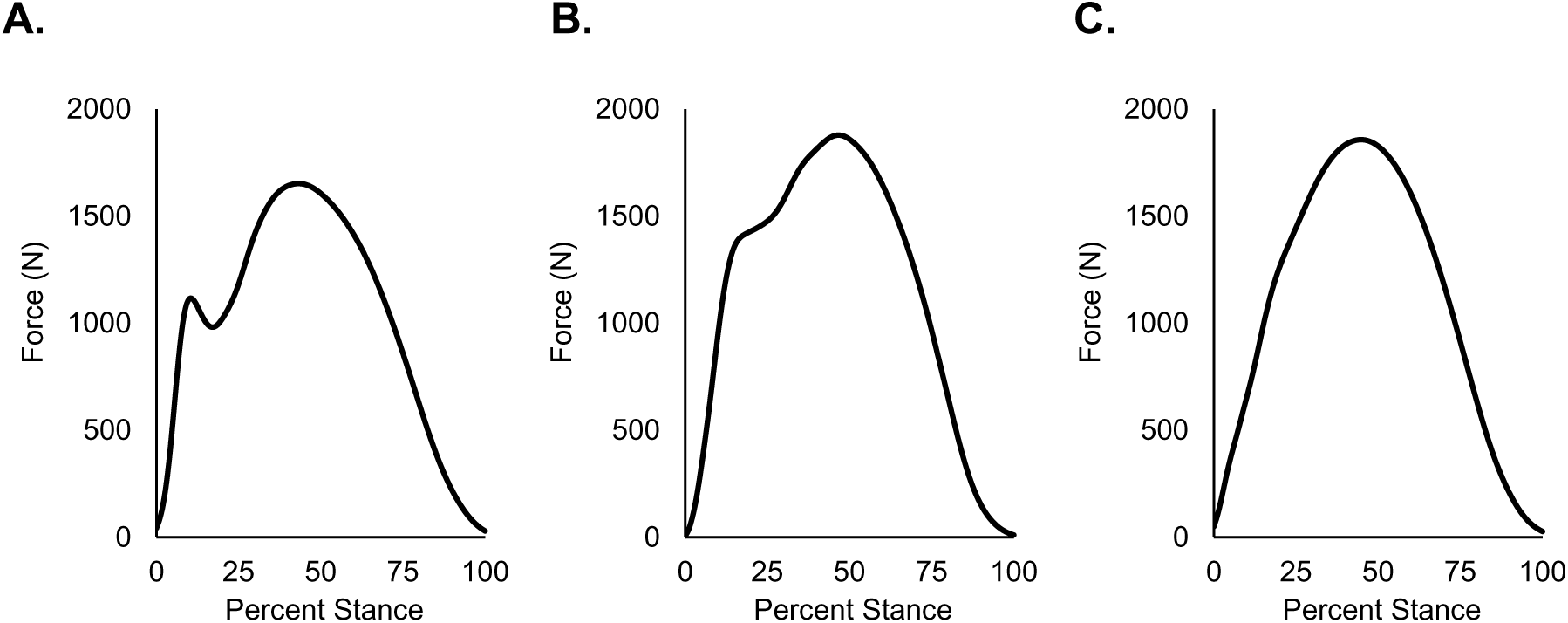
Exemplar vertical ground reaction forces. A prominent impact force tends to indicate rearfoot pattern (A); blunted impact force tends to indicate a midfoot pattern (B); and visually absent impact force tends to indicate a forefoot pattern (C).

**Table 1:** Participant characteristics. Mean ± SD participant characteristics of the rearfoot group and the forefoot group. d_MT_ is the standing Achilles tendon moment arm measured using the methods of Scholz et al. (2008). There were no statistically significant differences between groups (Student’s t-test, α = 0.05).

| | Males/<br>Females<br>(#) | Age<br>(years) | Height<br>(m) | Mass<br>(kg) | Pref. Speed<br>(m•s<sup>-1</sup>) | km/week<br>(km) | $d_{MT}$<br>(cm) |
| --- | --- | --- | --- | --- | --- | --- | --- |
| Rearfoot | 12/5 | 27.1 ± 6.4 | 1.76 ± 0.08 | 70.5 ± 8.6 | 3.7 ± 0.3 | 47.9 ± 34.8 | 4.64 ± 0.67 |
| Forefoot | 14/3 | 24.8 ± 6.4 | 1.78 ± 0.08 | 70.8 ± 9.0 | 3.8 ± 0.3 | 49.2 ± 25.8 | 4.47 ± 0.26 |
| *p*-value | - | 0.301 | 0.637 | 0.916 | 0.735 | 0.904 | 0.338 |

### Protocol

Participants practiced both footfall patterns until they felt comfortable with the protocol. Participants then performed trials of over-ground running with both rearfoot and forefoot conditions, contacting the force platform with their right foot. Regardless of their habitual footfall pattern, all participants were instructed to “run by landing on your heel first” for the rearfoot condition and for the forefoot condition, to “land with your toes first while trying to keep your heel from touching the ground.” Trials were retained for analysis if the average speed between the photogates was within ±5% of 3.5 m•s^−1^ (3.33-3.68 m/s), if the participant exhibited the proper footfall pattern using the previously noted criteria, and if the participant did not visibly alter their gait mechanics to contact the force platform. A minimum of ten trials were retained for each condition and participant.

Visual 3D software (C-Motion, Rockville, MD, USA) was used to measure leg length from the reflective markers on the lateral malleoli to the lateral femoral condyle of the right leg during the standing calibration trial. A high-resolution digital camera (Exilim EX-F1, Casio Computer Co., LTD, Shibuya-ku, Tokyo, Japan) was used to photograph the location of the lateral malleolus, and its center was marked with a pen. The Achilles tendon moment arm was measured from the center of the lateral malleolus to the posterior aspect of the Achilles tendon (Scholz et al., 2008).

### Data Analysis

The order of footfall pattern conditions was randomized between participants and counterbalanced between groups. Visual3D software was used to calculate 3D ankle joint angle, knee joint angle, and internal ankle joint moment from the motion capture and GRF data (Hamill et al., 2014; Selbie et al., 2014). Time series of these variables were linearly interpolated to 101 points per ground stance phase then averaged over trials for each participant. These data served as the input for the musculoskeletal model (Figure 2).

**Figure 2:**
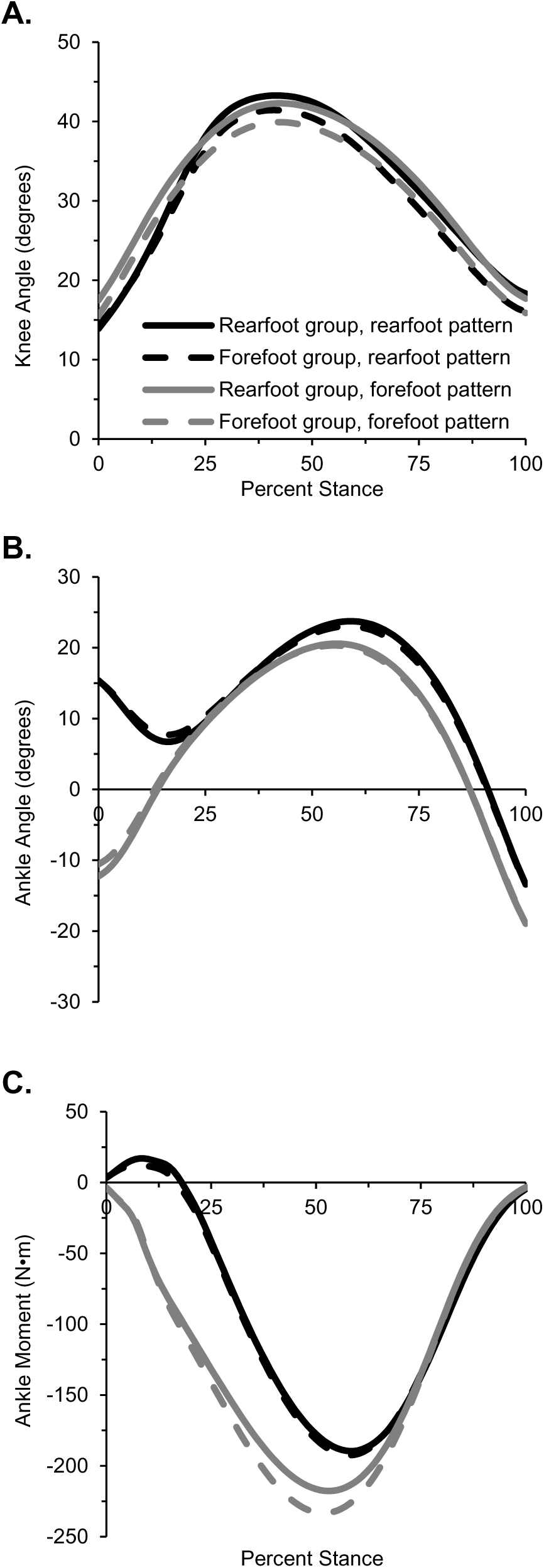
Model input variables. Group mean (A) knee joint angle, (B) ankle joint angle, and (C) ankle joint moment during the stance phase of rearfoot (solid) and forefoot (dashed) pattern performed by the rearfoot (black, n=17) and forefoot (grey, n=17) groups.

Kinovea Motion Tuner software v. 0.8.15 (www.kinovea.org) was used to calculate the static Achilles tendon moment arm length from the digital photograph similar to methods reported previously (Scholz et al., 2008). The static Achilles tendon moment arm was defined as the Euclidean distance from the line of action of the Achilles tendon (posterior aspect over the skin) to the center of rotation of the ankle. The center of rotation of the ankle was approximated as the midpoint between the medial and lateral malleoli (Lundberg et al., 1989). There was no statistical difference in static Achilles tendon moment arm length observed between groups (Table 1).

### Musculoskeletal Model

A list of abbreviations and acronyms is provided in Table 2. A two-dimensional (sagittal plane) musculoskeletal model of the thigh, shank, and foot (Figure 3) was developed in a MATLAB computer program (MathWorks, Natick, MA, USA). Details and specific equations about the model are described in Supplemental Material A but are explained in brief here. The model had two degrees of freedom (knee and ankle joint angular positions) and two muscles representing the triceps surae. Muscle moment arms were defined as second-order polynomial functions of the joint angles using data from the model of Arnold et al. (2010) with the soleus and gastrocnemius moment arms in the neutral ankle position defined on a participant-specific basis from the measured Achilles tendon moment arms. Equations for muscle-tendon unit (MT) origin-to-insertion lengths were then derived from integration of the moment arm equations using the virtual work principle (An et al., 1984). Neutral MT lengths were referenced from Arnold et al. (2010). The zero-order coefficient of polynomial equations for the muscle moment arm and MT were scaled for each participant individually by the static Achilles tendon moment arm and leg length, respectively. Equations for MT origin-to-insertion velocity were derived by symbolic differentiation of the muscle length equations with respect to time.

**Figure 3:**
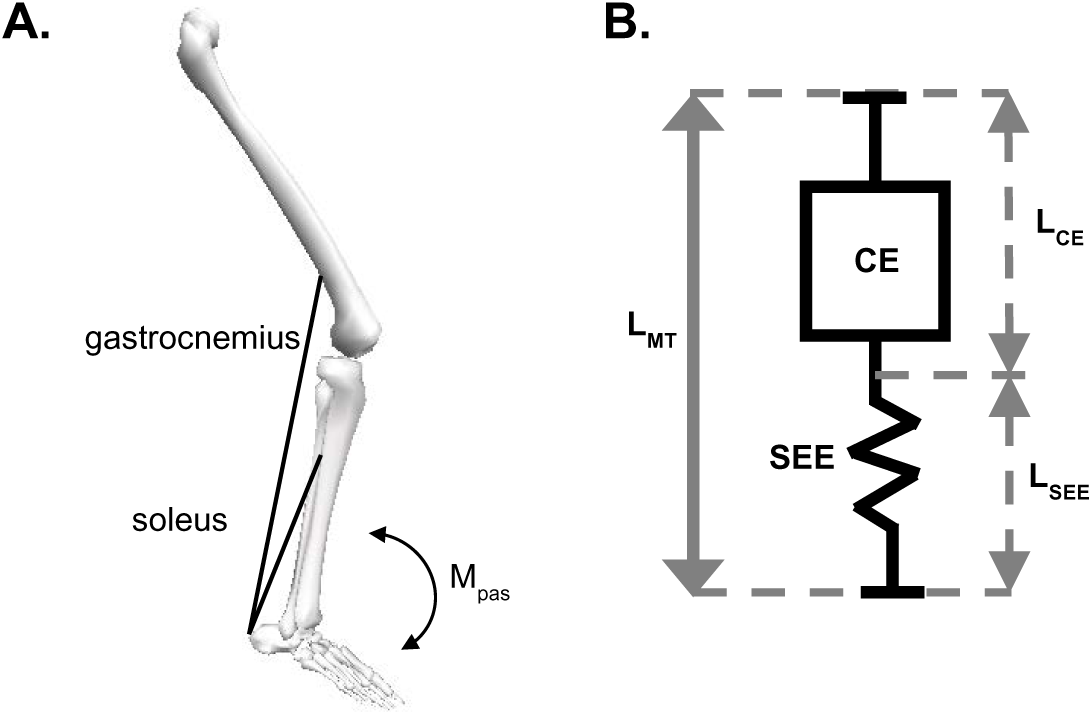
Musculoskeletal model. A) Link segment model representing the foot, leg, thigh, gastrocnemius, soleus, and passive moment (M_pas_) (Adapted from Nagano et al., 2001). B) Elements of the Hill-type model representing each muscle. The elements include a contractile element (CE) and series elastic element (SEE). The lengths of the CE (L_CE_) and SEE (L_SEE_) were constrained so the sum was equal to the muscle-tendon length (L_MT_).

**Table 2:** Acronyms and abbreviations.

|  |  |  |  |
| --- | --- | --- | --- |
| CE | contractile element | $L_{SEE}$ | series elastic element length |
| $CE_{MEE}$ | CE metabolic energy expenditure | $M_{pas}$ | passive moment |
| $d_{MT}$ | measured Achilles tendon moment arm length | MT | muscle-tendon complex |
| GRF | ground reaction force | SEE | series elastic element |
| $L_0 \cdot s^{-1}$ | optimal CE lengths per second | $V_{MAX}$ | maximum contractile element contraction velocity |
| $L_{CE}$ | contractile element length | $U_{MAX}$ | relative elongation of the SEE at maximum isometric force production |
| $L_{MT}$ | muscle-tendon complex length | | |

Each muscle was a Hill-based model with a contractile element (CE) in series with an elastic element (SEE). Parallel elasticity was implemented at the joint level according to Riener & Edrich (1999). The muscle model CE-SEE contractile dynamics consisted of the CE force-length relationship and SEE force-extension relationship from van Soest & Bobbert (1993) and CE force-velocity relationship from Epstein & Herzog (1998). Muscle-specific model parameters were derived from the literature (Supplemental Material A, Table S1). The internal states of the muscle model were based on the experimental data and constrained by the muscle geometry of the equilibrium condition: the length of the MT equaled the sum of the CE and SEE lengths, and the force generated in the muscle was equal in the MT, CE, and SEE.

At each 1% of the stance phase, the muscle moment arms were calculated from the joint angle inputs. The active ankle moment was calculated by subtracting the passive moment from the sagittal plane inverse dynamics moment (Riener and Edrich, 1999). Gastrocnemius and soleus muscle forces were calculated from the active moment assuming the moment arm lengths were force-dependent, increasing in length as force increases (Maganaris et al., 1998), and assuming a soleus/gastrocnemius force ratio based on the physiological cross-sectional areas scaled by body mass (Handsfield et al., 2014). Gastrocnemius and soleus forces were set to zero if the active ankle moment was a dorsiflexor moment, which only occurred in both groups within the first 18% of stance during rearfoot running on average across participants and trials. The SEE force was input to the force-extension relationship to calculate the SEE length. SEE length was subtracted from the MT length to calculate the CE length. MT and SEE velocity were calculated by differentiating SEE length with respect to time. CE velocity was calculated by subtracting SEE velocity from MT velocity.

Muscle activation was calculated from the contraction dynamics of the CE based on the Hill muscle model force-length and force-velocity relationships (Epstein and Herzog, 1998; van Soest and Bobbert, 1993). Muscle activation level was calculated at each 1% of the stance phase given the instantaneous CE length, CE velocity, and CE force. A reserve torque was calculated for time steps that resulted in activations above 1.0 for either muscle (Hicks et al., 2015) (Supplemental Material A). The reserve torque was invoked during the early and late portions of the stance phase, representing when the triceps surae first became activated in early stance and when the muscles were experiencing rapid shortening velocities in late stance (Figure S-A2). During rearfoot running, the reserve torque was invoked for a total of 182 data points across the stance phase of all participants. That is, the reserve torque was invoked for 5% of the total 3636 data points across participants during rearfoot running; 2.0% occurred between 0%–29% of the stance phase and the other 3.0% occurred between 82%–100% of the stance phase. The reserve torque was invoked for a total of 399 data points across the stance phase of forefoot running across all participants (Figure S-A2). That is, the reserve torque was invoked for approximately 11% of the total 363 data points across all participants during forefoot running; 2.6% occurring between 0%–41% of the stance phase and the other 8.4% occurring between 77%–100% of the stance phase.

The mechanical power developed by the MT, CE, and SEE was calculated as the product of the MT, CE, and SEE force and the respective MT, CE, and SEE velocity for each instant of the stance phase. Positive mechanical work (indicating energy generation) and negative mechanical work (indicating energy absorption) of the MT, CE, and SEE were then calculated by trapezoidal integration of the positive and negative potions of the power-time series, respectively. Net mechanical work was the sum of the positive and negative work occurring within the stance phase. The elastic strain energy stored and returned by the tendon and other elastic structures of the muscle during stance phase was defined as the amount of negative and positive mechanical work performed by the SEE, respectively.

The CE metabolic power was calculated as a function of activation, CE length, and CE velocity using the energetics model of Minetti & Alexander (1997) transformed to linear velocity variables (Sellers et al., 2003). CE metabolic energy expenditure (CE_MEE_) was calculated across the entire stance phase by integrating metabolic power with respect to time.

### Sensitivity Analysis

It is unknown currently if the muscle model parameter values should differ between habitual rearfoot and forefoot runners. Therefore, a Monte Carlo sensitivity analysis was performed to assess the effects of the chosen parameter values for tendon slack length, optimal fiber length, and muscle volume on the model outputs (Miller et al., 2017). The mean ±1 standard deviation of the participant-specific values of these parameters were used to determine the standard normal distribution from which new values were randomly selected. New parameter values were recalculated for each participant, the model simulation was performed based on new parameter values for all participants, and then the statistical analysis on the model output was repeated. The process continued until the fraction of iterations with a significant difference between groups or footfall patterns changed by ≤1% over 100 further iterations. An additional parameter sweep was performed to assess if the statistical results were affected by the values selected for (1) the relative elongation of the SEE at maximum isometric force production (U_MAX_) and for (2) the maximum CE shortening velocity (V_MAX_). Values tested for U_MAX_ ranged from 0.03–0.12 in increments of 0.01. Values tested for V_MAX_ ranged from 9–16 optimal CE lengths per second (L_0_•s^−1^) in increments of 1.0. A third sensitivity analysis was performed for which maximum isometric force generation was increased for participants in the forefoot group based on the differences in isometric plantar flexor maximum voluntary contraction between rearfoot and forefoot runners reported by Liebl et al. (2014).

### Statistical Analysis

Peak activation, peak muscle force, mechanical work of the MT, CE, and SEE, and CE_MEE_ were compared between the rearfoot and forefoot running patterns and groups across the stance phase. Each outcome variable was subjected to a mixed model ANOVA with footfall pattern and group as fixed variables and participant nested within group as a random variable. The differences between footfall patterns (rearfoot and forefoot), between groups (habitual rearfoot and habitual forefoot runners), and the interaction of footfall pattern and group were assessed with a significance level of α = 0.05. The ANOVA results were supplemented by effect size (*d*) calculations to assess substantive meaningfulness of differences between comparisons. Effect size was calculated as the difference between the means divided by the pooled standard deviation. An effect size of less than 0.5 indicated a small effect, an effect size between 0.5 and 0.7 indicated a moderate effect, and an effect size greater than 0.8 indicated a large effect (Cohen, 1992).

## Results

Primary results are presented here, while more detailed kinematic, kinetic, and activation results for the soleus and gastrocnemius MTU, CE, and SEE during rearfoot and forefoot running are presented in Supplemental Material B. Given that there were no significant interactions, the following results for mechanical work variables and CE_MEE_ were collapsed across groups because only footfall pattern main effects were observed for all statistically significant comparisons.

### Muscle activation and force

Peak gastrocnemius activation was 28.3% greater during rearfoot running compared with forefoot running (*P*<0.001, *d*=1.34) and occurred at approximately 25% of the stance phase during both footfall patterns (Figure 4A). However, gastrocnemius activation was greater during forefoot running than during rearfoot running until approximately 75% of the stance phase (Figure 4A). Soleus activation was greater during forefoot running throughout the stance phase compared with rearfoot running (Figure 4D). Peak soleus activation was not different between footfall patterns (*P*>0.05, *d*<0.25) but soleus activation throughout the stance phase was greater during forefoot running compared with rearfoot running (Figure 4D). MT force of both muscles was greater in forefoot running than during rearfoot running from 0%–73% of the stance phase (Figure 4B&E). Peak muscle force was produced at 58% and 53% of stance during rearfoot and forefoot running, respectively, in both muscles. Peak gastrocnemius and soleus force was 17.6% greater during forefoot running than rearfoot running (gastrocnemius: *P*<0.001, *d*=1.00; soleus: *P*<0.001, *d*=0.92) (Figure 4B&E). In the first half of the stance phase, gastrocnemius CE was concentric primarily during both footfall patterns, but the shortening velocity was slower during forefoot running than rearfoot running (Figure 4C). Peak CE shortening velocity in the gastrocnemius was similar between footfall patterns during late stance (*P*>0.05, *d*<0.10). The soleus acted primarily eccentrically until approximately 50% of the stance phase of forefoot running, whereas the soleus acted concentrically during this period of stance during rearfoot running (Figure 4F). Peak CE shortening velocity in the soleus was 7.3% greater during late stance in forefoot running compared with rearfoot running (*P*<0.001, *d*=0.33) (Figure 4F).

**Figure 4:**
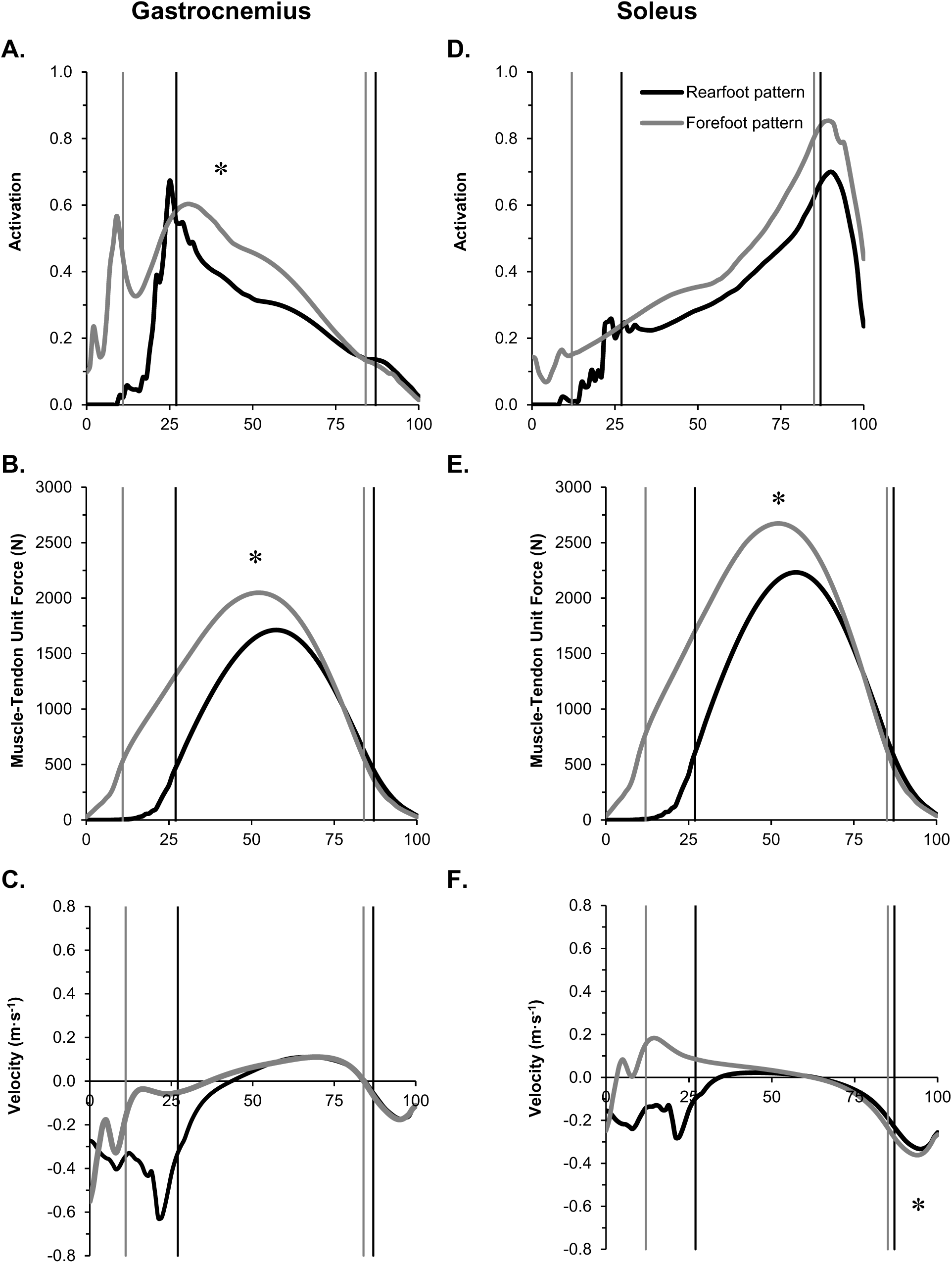
Muscle activation, muscle force generation, and contractile element velocity. Mean activation, muscle-tendon complex force, and contractile element velocity during the stance phase of rearfoot (black) and forefoot (grey) pattern running, collapsed across the rearfoot and forefoot groups (n=34). Data for the gastrocnemius are presented in panels A–C. Data for the soleus are presented in panels D–F. Vertical lines in each panel indicate the range of time when the muscle was generating greater than 25% of the peak force produced during the stance phase. * indicates a statistically significant difference between footfall patterns (*P*<0.05).

### Muscle mechanical work

Significant footfall pattern main effects were observed for all mechanical work variables of the gastrocnemius and soleus (*P*<0.010, *d*=0.37–3.49) except for positive CE work by the gastrocnemius, which was not different between footfall patterns (*P*>0.05, *d*=0.18). Net SEE work by the gastrocnemius as well as net work of the MT, CE, and SEE by the soleus were positive during rearfoot running and negative during forefoot running (Figure 5A,D). Net MT and CE mechanical work by the gastrocnemius were both negative and greater during forefoot running than rearfoot running (Figure 5A). For both muscles, forefoot running resulted in greater positive and negative work of the MT and SEE than rearfoot running (Figure 5B,C,E,F). Forefoot running also resulted in greater negative CE work by the gastrocnemius as well as positive and negative CE work by the soleus compared with rearfoot running. Effect sizes were large (i.e., *d*=0.84 – 3.49) for all mechanical work variables for both muscles except for positive CE work (gastrocnemius *d*=0.18; soleus *d*=0.37), net SEE work (gastrocnemius *d*=0.41; soleus *d*=0.38), and positive MT work (gastrocnemius *d*=0.52; soleus *d*=0.65), as well as negative CE work by the gastrocnemius (*d*=0.65).

**Figure 5:**
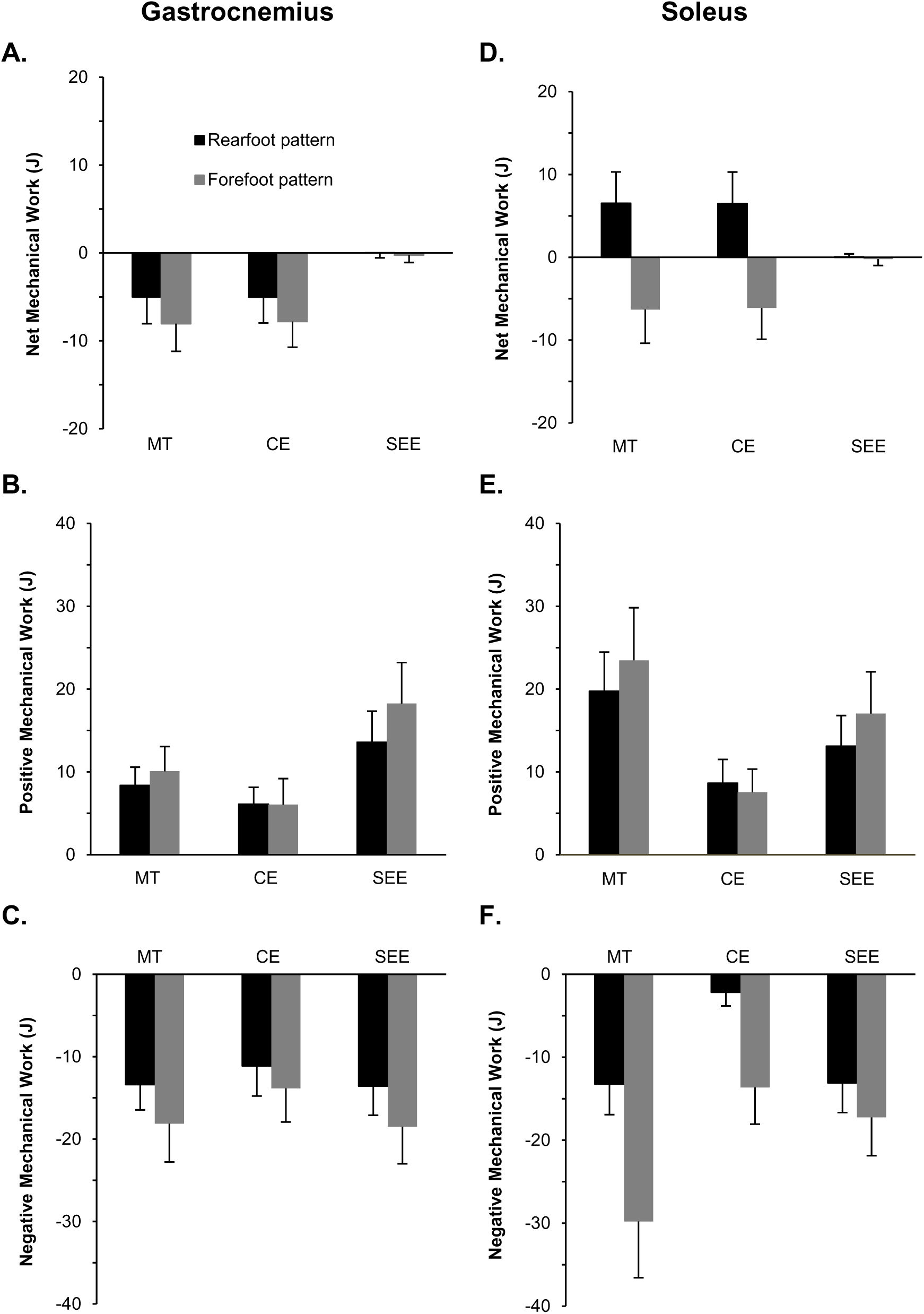
Muscle element mechanical work production. Mean mechanical work produced by the muscle-tendon complex (MT), the contractile element (CE), and the series elastic element (SEE) of the gastrocnemius (panels A–C) and soleus (panels D–F) during the stance phase of rearfoot (black bars) and forefoot (grey bars) pattern running. Data are collapsed across the rearfoot and forefoot groups (n=34). Error bars are ±1SD. All mechanical work variables of the gastrocnemius and soleus were significantly different (*P*<0.010) except for CE positive work done by the gastrocnemius (*P*>0.05).

### Muscle metabolic energy expenditure

Differences in gastrocnemius CE metabolic power between groups and footfall patterns occurred before ~50% of the stance phase and was nearly identical between footfall patterns for the remainder of stance (Figure 6A). In the soleus, CE metabolic power was similar between footfall patterns until ~60% of the stance phase (Figure 6B). Forefoot running resulted in 3.5% greater gastrocnemius CE_MEE_ compared with rearfoot running but this difference was not significant between footfall patterns (*P*>0.05, *d*=0.08; Figure 6C). Soleus CE_MEE_ was 12.2% greater during forefoot running compared with rearfoot running (Figure 6C). Although this difference was significant (*P*<0.001), only a small effect size was observed (*d*=0.34).

**Figure 6:**
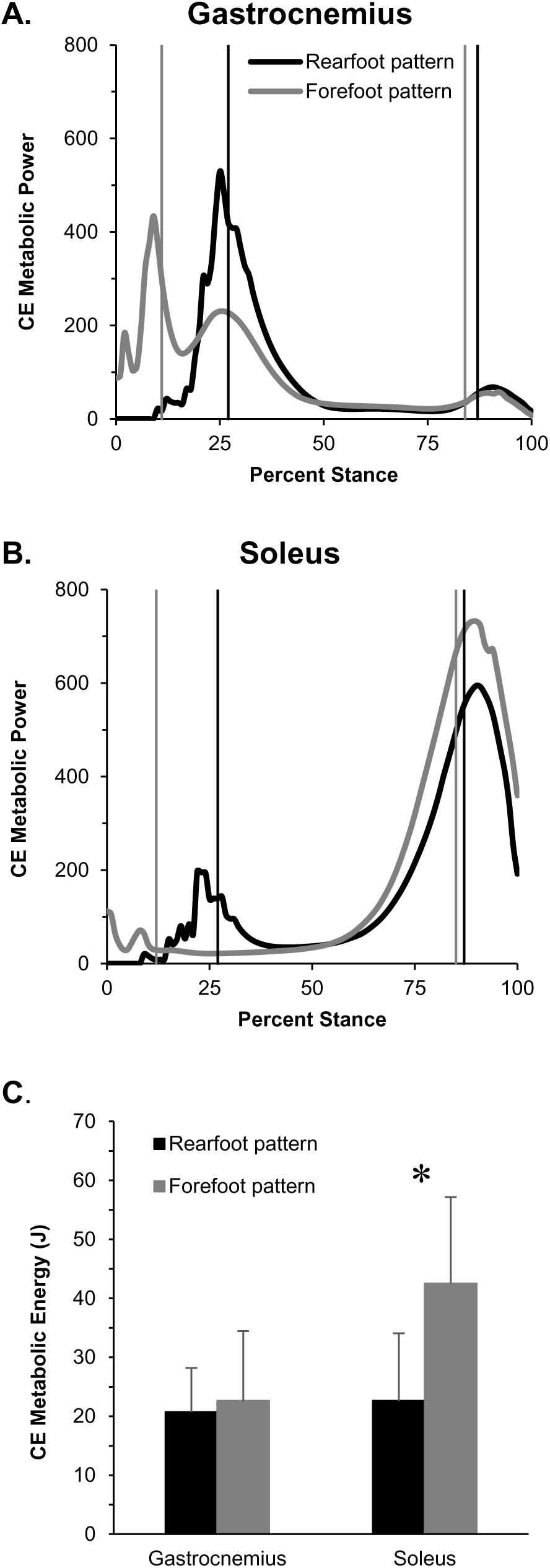
Contractile element metabolic power and metabolic energy expenditure. Mean contractile element (CE) metabolic power of the gastrocnemius (panel A) and soleus (panel B) during the stance phase of rearfoot (black line) and forefoot (grey line) pattern running, collapsed across the rearfoot and forefoot groups (n=34). Vertical lines indicate the range during stance when the muscle was generating greater than 25% of the peak force produced. (C) Mean CE metabolic energy expenditure produced during stance of rearfoot (black bars) and forefoot (grey bars) running in the gastrocnemius and soleus muscles. Error bars are ±1SD. * indicates a statistically significant difference between footfall patterns (*P*<0.05).

### Sensitivity analysis

#### Monte Carlo Sensitivity Analysis

The Monte Carlo sensitivity analysis on tendon slack length, optimal fiber length, and muscle volume was performed with the model results collapsed across groups given that only footfall pattern main effects were found. The Monte Carlo sensitivity analysis converged after approximately 640 iterations (Figure 7). Net MT work by the soleus, negative MT work by the soleus, net CE work by the soleus, and negative CE work by the soleus were statistically different between forefoot running and rearfoot running for 87.8%, 91.0%, 99.0%, and 99.3% of these iterations, respectively (*P*<0.05). No other mechanical work variables were statistically different for more than 50% of these iterations (*P*>0.05).

**Figure 7:**
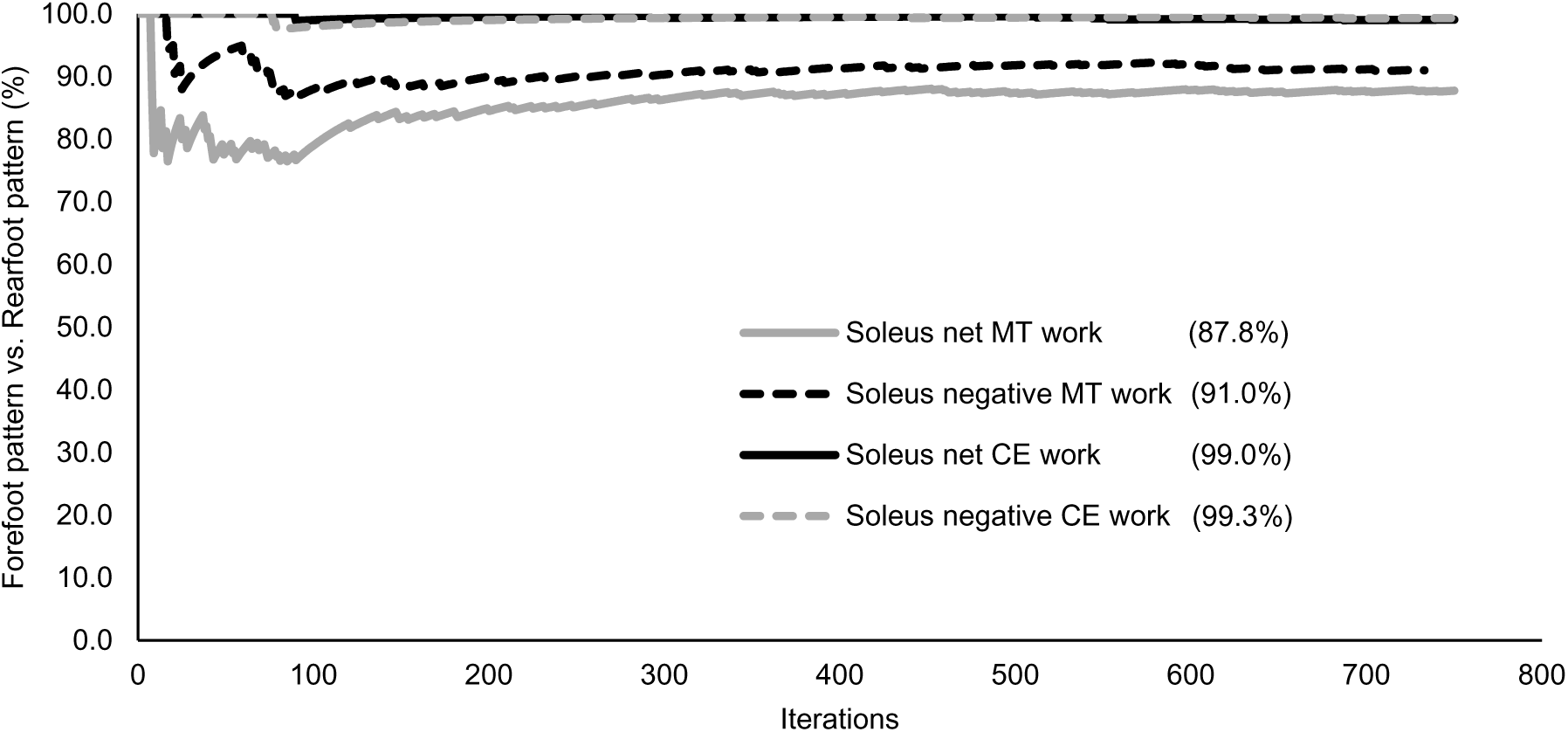
Monte Carol sensitivity analysis running fraction. Vertical axis is the fraction of iterations for which forefoot pattern running resulted in greater net contractile element (CE) work, net muscle-tendon complex (MT) work, and negative CE work of the soleus compared with rearfoot pattern running (*P*<0.05). Data are collapsed across the rearfoot and forefoot groups (n=34). Horizontal axis is the number of iterations performed. Net MT work by the soleus, net CE work by the soleus, and negative CE work by the soleus remained statistically greater during forefoot running than during rearfoot running for 93.2%, 100%, 86.6% of the iterations, respectively (*P*<0.05). Therefore, the statistical results for these mechanical work variables of the soleus were not dependent on the model parameter values selected for tendon slack length, optimal fiber length, and muscle volume.

#### Sensitivity to U_MAX_

Changing U_MAX_ of the gastrocnemius affected the statistical relationship between footfall patterns for only positive CE work, net SEE work, positive MT work, and CE_MEE_. Rearfoot running resulted in significantly greater positive CE work by the gastrocnemius for U_MAX_ values 0.03–0.09 compared with forefoot running (*P*<0.021; d=0.34–1.06). The difference in net SEE work by the gastrocnemius between footfall patterns became nonsignificant at U_MAX_ values 0.11–0.12 (*P*>0.05; d=0.09–0.33). The difference in positive MT work by the gastrocnemius between footfall patterns became nonsignificant for U_MAX_ values 0.03 – 0.04 (*P*>0.05; d = 0.09–0.17). Gastrocnemius CE_MEE_ became statistically greater in rearfoot versus forefoot running for U_MAX_ values 0.03–0.08 (*P*<0.004; d=0.27–0.81) but statistically smaller during rearfoot running for U_MAX_ values 0.11–0.12 (*P*<0.046; d=0.31–0.47).

For the soleus, changing U_MAX_ affected the statistical relationship between footfall patterns for only positive CE work by the soleus for U_MAX_ values below 0.08, as well as net SEE work by the soleus and soleus CE_MEE_ for U_MAX_ values 0.11 or greater. Specifically, the difference between footfall patterns in positive CE work by the soleus became nonsignificant for U_MAX_ values 0.03–0.08 (*P*>0.05; d=0.09–0.16). The difference in net SEER Rwork by the soleus and soleus CE_MEE_ between footfall patterns became nonsignificant for U_MAX_ values 0.11–0.12 (*P*>0.05; d=0.10–0.34) and 0.12, respectively (*P*>0.05; d=0.15).

#### Sensitivity to V_MAX_

Changing V_MAX_ affected the statistical results for only net SEE work and CE_MEE_ of both muscles. In the gastrocnemius and soleus, the difference in net SEE work between footfall patterns became nonsignificant for V_MAX_ values 14–16 L_0_•s^−1^ (*P*>0.05, d=0.06–0.33). Gastrocnemius CE_MEE_ became statistically greater during forefoot running compared with rearfoot running when V_MAX_ was 9 L_0_•s^−1^ (*P*=0.041, d=0.40). The difference in soleus CE_MEE_ between footfall patterns became nonsignificant when V_MAX_ was 9 and 10 L_0_•s^−1^ (*P*>0.05, d=0.06–0.16).

#### Sensitivity to F_MAX_

Increasing the gastrocnemius and soleus maximum isometric force capability of the forefoot group did not affect the direction of the results or the statistical outcome for any muscle mechanical work or muscle metabolic energy expenditure variable.

## Discussion

The purpose of the present study was to compare the muscle mechanical work and predicted muscle metabolic energy expenditure of the gastrocnemius and soleus between rearfoot and forefoot running patterns. The first hypothesis was that rearfoot running would result in the gastrocnemius and soleus producing greater CE mechanical work than forefoot running, whereas forefoot running would result in greater SEE mechanical work (i.e., greater storage and release of elastic energy) from the gastrocnemius and soleus compared with rearfoot running. This hypothesis was not supported with respect to CE mechanical work given that net and negative CE work in both muscles was greater during forefoot running compared with rearfoot running (Figure 5). Rearfoot running resulted in greater positive CE work than forefoot running in both muscles, but the difference was not statistically significant in the gastrocnemius. The first hypothesis was supported with respect to the SEE in both muscles because forefoot running resulted in greater SEE work than rearfoot running (Figure 5). Therefore, storage and release of elastic energy was greater during forefoot running, but it was not effective at reducing CE mechanical work compared with rearfoot running.

The results for SEE mechanical work support the previous suggestion that forefoot running results in a greater contribution from elastic energy compared with rearfoot running (Ardigo et al., 1995; Hasegawa et al., 2007; Lieberman et al., 2010; Perl et al., 2012). However, the second hypothesis, that forefoot running would result in lower triceps surae metabolic energy expenditure than rearfoot running, was not supported. Greater SEE mechanical work production will only affect muscle metabolic energy expenditure if there is also less CE mechanical work. However, forefoot running resulted in greater positive and negative CE mechanical work by the soleus and greater negative CE mechanical work by the gastrocnemius than rearfoot running (Figure 5). Greater CE mechanical work during forefoot running, despite greater elastic energy contribution (i.e., SEE mechanical work), may partially explain why forefoot running resulted in similar gastrocnemius metabolic energy expenditure and greater soleus metabolic energy expenditure than rearfoot running (Figure 6C). In the gastrocnemius, greater net mechanical work was done during forefoot running but at the same metabolic cost as rearfoot running. In the soleus, greater overall CE work was done without a comparable increase in SEE work during forefoot running than rearfoot running. Therefore, greater SEE mechanical work enhanced mechanical work production of the muscle-tendon unit in the gastrocnemius and soleus during forefoot running, but it did not result in reduced muscle metabolic energy expenditure than rearfoot running.

The similarity in metabolic energy expenditure of the gastrocnemius between rearfoot and forefoot running may be a result of the offsetting mechanical and metabolic effects of CE shortening velocity and force generation. Forefoot running required greater activation in order to generate greater gastrocnemius force, but this force was produced more economically because of slower CE contraction velocities compared with rearfoot running. In the gastrocnemius, rearfoot running resulted in greater CE shortening velocity until ~35% of stance, less force production until ~70% of stance, and less activation until ~75% of stance compared with forefoot running (Figure 4A–C). The faster CE shortening velocity by the gastrocnemius during rearfoot running resulted in a reduced force generation capability and greater metabolic cost than if CE shortening velocity by the gastrocnemius was slower or isometric. However, rearfoot running required less force production than forefoot, which kept activation lower than if a high CE shortening velocity and high force generation were required. Therefore, a lower force requirement and activation of rearfoot running resulted in similar gastrocnemius metabolic cost as forefoot running, despite slower CE shortening velocities during forefoot running.

Soleus CE velocity was nearly isometric for the majority of stance during rearfoot running, whereas forefoot running resulted in CE lengthening by the soleus for the first half of stance, followed by a brief isometric phase through mid-stance (Figure 4F). The primary differences in CE velocity occurred before muscle the muscle was generating over 25% of the peak force produced. These differences in CE velocity by the soleus caused more negative CE work to occur with forefoot running than rearfoot running. Therefore, more soleus work was dissipated by the CE during forefoot running than generated by the CE during either footfall pattern (Figure 5D-F). Although negative CE work carries a lower metabolic cost than positive (Biewener and Roberts, 2000), the differences in negative CE work by the soleus cannot explain the differences in soleus metabolic cost between footfall patterns given metabolic cost was similar when negative CE work by the soleus was being produced (Figure 6B). Soleus metabolic energy expenditure did not differ between footfall patterns until approximately 65% of stance because of offsetting metabolic effects of contraction velocity and force production: Rearfoot running resulted in faster CE shortening velocities by the soleus but lower activation and force production than forefoot running during this period of stance (Figure 4D-F). During push-off, however, forefoot running resulted in greater CE work by the soleus and CE metabolic energy expenditure as a result of greater activation and shortening velocity compared with rearfoot running. This greater activation and CE metabolic energy expenditure occurred despite soleus force production being similar between patterns during the last third of the stance phase.

There were no significant group main effects in mechanical or metabolic work of the gastrocnemius or soleus between habitual rearfoot and forefoot runners (Figures 5 and 6). Therefore, the muscle dynamics of each footfall pattern, and the associated metabolic cost, was likely not a result of training with or being habituated to a given footfall pattern. Liebl et al. (2014) recently found that habitual forefoot runners could generate 28% greater isometric plantar flexor force during a maximum voluntary contraction than habitual rearfoot runners. This result suggests that habitually running with a forefoot pattern results in muscular strength adaptations to the plantar flexors. However, increasing maximum isometric force generation in the muscle model of the present study did not result in group main effects, or influence the muscle dynamics or metabolic cost of the gastrocnemius or soleus between footfall patterns. Furthermore, the results obtained during the parameter sweep sensitivity analysis were consistent with the direction of the results from the original model. The statistical results of the parameter sweep did not differ from the original model, except for some cases toward the extremes of the tested parameters. Therefore, the differences in muscle dynamics and muscle metabolic cost of the triceps surae muscles between footfall patterns are likely not affected by modest gains in muscle force generation capacity or choice of modeling parameter values.

Consistent with the results of the present study, authors of previous studies suggested that greater force transmission through the Achilles tendon during forefoot running would result in greater elastic energy storage compared with rearfoot running (Almonroeder et al., 2013; Ardigo et al., 1995; Hasegawa et al., 2007; Nilsson and Thorstensson, 1989; Perl et al., 2012). Ardigo et al. (1995) indirectly estimated the contribution of elastic energy between footfall patterns by calculating the ratio between external work and deceleration time to external work and acceleration time. These authors found that forefoot running resulted in greater estimated elastic energy storage compared with rearfoot running, although no difference in sub-maximal oxygen consumption was observed between footfall patterns. These authors suggested that greater storage of elastic energy during forefoot running compensated for the additional external work observed with this pattern compared to rearfoot. Additionally, Hof et al. (2002) reported that the triceps surae of an individual midfoot runner produced less CE work than that of an individual rearfoot runner. Some individuals in the present study showed the same response as found by Hof et al. (2002), but on the group level in the present study, we found that forefoot running resulted in greater CE work than rearfoot running. Between-subject variation likely explains the differences in results between Hof et al. (2002) and the present study, in addition to the possible, but likely minor, differences in muscle mechanics between midfoot verses forefoot running. The present study expands on these earlier results by demonstrating that greater activation and force generation required with forefoot running offset any potential metabolic savings that could result from greater storage and release of elastic energy compared with rearfoot running.

A study by Heise et al. (2011) found that more economical runners tended to perform less negative work at the ankle. The present study was consistent with these results given that rearfoot running resulted in smaller negative work of the triceps surae muscle-tendon complex (Figure 5C&F) and also resulted in a smaller predicted metabolic cost of the soleus compared with forefoot running (Figure 6). Although the ankle plantar flexors produce greater positive joint work during running compared with the muscles of the hip and knee (DeVita et al., 2008; Heise et al., 2011; Stefanyshyn and Nigg, 1998; Winter, 1983), the triceps surae muscle-tendon complex has a relatively small active muscle volume during running than other muscles of the lower extremity. Therefore, the muscle-tendon interactions of larger muscles of the lower extremity may have a greater effect on the whole-body metabolic cost during running than the triceps surae. For example, the quadriceps undergoes greater SEE elongation and elastic energy release in more economical runners (Albracht and Arampatzis, 2006). Forefoot running results in a slightly less knee flexion angle at ground contact than rearfoot running, but the knee flexion angle is similar throughout the rest of stance (Figure 2A). Although the difference in elastic energy contribution or metabolic energy expenditure of the quadriceps between footfall patterns is currently unknown, the similarity of the knee joint angles during the stance phase overall suggests that muscle dynamics and metabolic cost of the quadriceps may also be similar between footfall patterns.

Additional sources of elastic strain energy or passive mechanical work during human locomotion have been identified. Series ankle elasticity, for example, can reduce total mechanical work during walking by redirecting the center of mass velocity and reducing energy loss due to the collision with the ground (Zelik et al., 2014). Soft tissue deformation and its subsequent rebound may also contribute 4 J of total positive work per stride of walking (Zelik and Kuo, 2010). The longitudinal arch of the foot can store approximately 17 J of elastic strain energy which is about half of the strain energy stored in the Achilles tendon under the same load (Ker et al., 1987). Together, the Achilles tendon and longitudinal arch of the foot can store approximately 50% of the mechanical energy stored in the body during the stance phase (Alexander and Bennet-Clark, 1977; Ker et al., 1987). It was recently reported that barefoot forefoot running resulted in greater estimated longitudinal arch strain during the stance phase compared with barefoot rearfoot running (Perl et al., 2012). The authors concluded that more arch strain during forefoot running contributes to a reduced whole body metabolic cost compared with rearfoot running although whole body metabolic cost in that study was similar between footfall patterns. The present study demonstrated that greater elastic energy storage and release does not necessarily result in reduced metabolic cost. Given that other recent studies have not found a difference in submaximal rates of oxygen consumption between footfall patterns (Ardigo et al., 1995; Cunningham et al., 2010; Perl et al., 2012) or have found rearfoot running to be more economical (Gruber et al., 2013; Ogueta-Alday et al., 2013), it is unlikely that greater longitudinal arch strain or other sources of elastic strain energy occurring with forefoot running result in it being a more economical footfall pattern than rearfoot running.

There are several possible limitations of the present study. The present study assumed no co-contraction between the ankle plantar flexor and dorsiflexor muscles. This assumption may have resulted in an underestimation of the amount of negative muscle-tendon complex work production by the gastrocnemius and soleus given force production of these muscles may have been greater than what can be calculated from joint moments determined from inverse dynamics for a brief period early in the stance phase. Furthermore, allocating force generation between the gastrocnemius and soleus based on PCSA assumes that force is shared equally per unit of PCSA, although it is possible that force-sharing may differ from that assumption. Additionally, the present study did not consider the effect of pre-activation of the gastrocnemius and soleus. Activation during both rearfoot and forefoot running was set to zero at the moment of initial ground contact in the present study, which may result in an underestimation of muscle activation during both footfall patterns. Forefoot running resulted in more gastrocnemius activation during terminal swing than rearfoot running in a recent study (Yong et al., 2014). Given this result, including pre-activation in the present model would further increase the overall activation and metabolic cost during forefoot running, but it would likely not affect the direction of the results.

## Conclusion

Forefoot running has previously been suggested to enhance metabolic economy compared with rearfoot running, due to greater storage and release of elastic energy of the triceps surae muscle complex in forefoot running. Although we found that forefoot running did result in a greater contribution of elastic energy to mechanical work of the gastrocnemius and soleus, predicted muscle metabolic energy expenditure by the soleus was greater in forefoot running compared with rearfoot running. These findings can be understood largely in terms of the CE force-velocity relation: a greater force or greater shortening velocity will favor greater mechanical work, but will also tend to have a greater metabolic cost due to greater muscle activation. In the gastrocnemius, force and velocity offset each other in forefoot versus rearfoot running; but greater activation and shortening velocity during push-off in the soleus resulted in greater soleus metabolic energy expenditure in forefoot running than rearfoot running. These results suggest that there is no muscle metabolic expenditure benefit of one footfall pattern over the other.

## Acknowledgments

The authors would like to thank the participants and Carl Jewell, Samuel del Pilar II, Dr. Lex Gidley for their contribution to this study.

